# Relationships between MA-RNA binding in cells and suppression of HIV-1 Gag mislocalization to intracellular membranes

**DOI:** 10.1101/629998

**Authors:** Dishari Thornhill, Balaji Olety, Akira Ono

**Author notes:** Address correspondence to Akira Ono, Ph.D. Balaji Olety, Ph.D. University of Massachusetts Medical School, 364 Plantation Street, LRB-526, Worcester MA 01605.

## Abstract

The HIV-1 Gag matrix (MA) domain mediates localization of Gag to the plasma membrane (PM), the site for infectious virion assembly. The MA highly basic region (HBR) interacts with phosphatidylinositol-(4,5)-bisphosphate [PI(4,5)P2], a PM-specific acidic lipid. MA-HBR also binds RNAs. To test whether acidic lipids alone determine PM-specific localization of Gag or whether MA-RNA binding also plays a role, we compared a panel of MA-HBR mutants that contain two types of substitutions at MA residues 25/26 or 29/31: Lys->Arg (KR) (25/26KR and 29/31KR) and Lys->Thr (KT) (25/26KT and 29/31KT). Consistent with the importance of the HBR charge in RNA binding, both KT mutants failed to bind RNA via MA efficiently unlike the corresponding KR mutants. Both 25/26KT Gag-YFP and 29/31KT Gag-YFP bound non-specifically to PM and intracellular membranes, presumably via the myristoyl moiety and remaining MA basic residues. In contrast, 25/26KR Gag-YFP bound specifically to the PM, suggesting a role for the total positive charge and/or MA-bound RNA in navigating Gag to the PM. Unlike 29/31KT Gag-YFP, 29/31KR Gag-YFP was predominantly cytosolic and showed little intracellular membrane binding despite having a higher HBR charge. Therefore, it is likely that MA-RNA binding blocks promiscuous Gag membrane binding in cells. Notably, introduction of a heterologous multimerization domain restored PI(4,5)P2-dependent PM-specific localization for 29/31KR Gag-YFP, suggesting that the blocking of PM binding is more readily reversed than that of intracellular membrane binding. Altogether, these cell-based data support a model in which MA-RNA binding ensures PM-specific localization of Gag via suppression of non-specific membrane binding.

**IMPORTANCE:** The PM-specific localization of HIV-1 Gag is a crucial early step in the infectious progeny production. The interaction between the MA highly basic region (HBR) of Gag and the PM-specific lipid PI(4,5)P2 is critical for Gag localization to the PM. Additionally, *in vitro* evidence has indicated that MA-RNA binding prevents non-specific binding of Gag to non-PI(4,5)P2-containing membranes. However, cell-based evidence supporting a role for HIV-1 MA-RNA binding in PM-specific subcellular localization has been scarce; thus, it remained possible that in cells, just the high basic charge or the PI(4,5)P2-binding ability is sufficient for MA to direct Gag specifically to the PM. The current study revealed for the first time an excellent correlation between RNA binding of MA-HBR and inhibition of promiscuous Gag localization, both within the cells, and thereby provided cell-based evidence supporting a mechanism in which HIV-1 MA binding to RNA ensures specific localization of Gag to the PM.

## Introduction

HIV-1 progeny virions exit most cells at the plasma membrane (PM) (1, 2). The matrix (MA) domain of the HIV-1 structural polyprotein Gag mediates the targeting and binding of Gag to the PM, which is a crucial stage of virus particle production (3–9). The two essential features of the MA domain required for the PM-specific localization and binding of Gag are an N-terminal myristate moiety and a highly basic region (HBR) (10–16). The myristoyl moiety allows Gag to form hydrophobic interactions with membranes when it is not sequestered in the MA globular domain (17–24). The MA-HBR comprises a highly conserved cluster of basic residues spanning residues 14-31 (13, 14, 23, 25). This basic patch drives electrostatic and head group-specific interactions of Gag with phosphatidylinositol (4,5)-bisphosphate [PI(4,5)P2], an acidic phospholipid found primarily in the inner leaflet of the PM (15, 20, 26–34). In addition to acidic phospholipids, RNA, which has been shown to bind the MA domain (35–43), may play an important role in regulation of Gag-membrane binding (28, 32, 44–48). Among the HBR basic residues, an NMR-based study showed that the residues 29 and 31 are particularly important for PI(4,5)P2 interaction in the absence of RNA (49).

WT Gag that is translated *in vitro* using rabbit reticulocyte lysates binds liposomes consisting of a neutral lipid, phosphatidylcholine (PC), and an acidic lipid, phosphatidylserine (PS) (PC+PS liposomes) poorly but shows enhanced membrane binding either when Gag is treated with RNase or when PI(4,5)P2 is included in the liposomes (32, 50). Furthermore, besides NC, MA-HBR mediates RNA binding to WT Gag in the cytosol, and removal of RNA enhances binding of such cytosolic Gag to PC+PS liposomes (46). In good agreement with these studies, RNase treatment of cell homogenates derived from HIV-1-expressing cells resulted in significant shift of Gag from cytosolic to membrane fraction (47). These observations suggest that WT Gag is susceptible to negative regulation of membrane binding by MA-bound RNA and that Gag-membrane binding occurs only when this RNA is removed by RNase or counteracted by PI(4,5)P2. Sequencing of RNAs cross-linked to MA revealed that the major RNA species bound to MA in cells is tRNA and that MA-tRNA binding is reduced with membrane-bound Gag compared to cytosolic Gag (47).

Based on these studies, our working model is that RNA bound to MA-HBR prevents Gag from electrostatically binding to acidic phospholipids such as PS, which are present ubiquitously in the cell (51). In this model, PI(4,5)P2 is able to overcome RNA-mediated negative regulation, thereby promoting Gag binding to the PM, while RNA prevents Gag from binding to other acidic lipids present in non-PM membranes(32, 44). The hypothesis that MA-RNA binding prevents promiscuous localization of Gag has not been directly investigated in the context of HIV-1 Gag expressed in cells. Our previous study of Gag chimeras containing various retroviral MA domains showed the presence of a correlation between the size of basic patches, RNA sensitivity in the *in vitro* liposome binding assay, and PM-specific Gag localization in cells (29). However, MA-RNA binding in cells was not measured in that study. Moreover, confounding effects of structural variations of the various retroviral MA domains, other than the size of the basic patches, cannot be excluded. In addition, although unlikely, it remains possible that Gag chimeras with different retroviral MA domains may be differentially endocytosed after nascent virion assembly at the PM, resulting in apparent differences in Gag localization.

In the current study, to examine the correlation between MA-RNA binding and subcellular Gag localization while addressing the limitations in the previous studies, we compared the effects of two types of amino acid substitutions in the context of HIV-1 MA-HBR and determined their ability to bind RNA in cells and the specificity of their localization. The first type is Lys-to-Thr (KT) changes. We previously showed that MA-HBR mutants with KT changes at MA residues 25 and 26 (25/26KT) or MA residues 29 and 31 (29/31KT) display promiscuous subcellular localization to both PM and intracellular membranes (14, 32). As for MA-RNA binding, previous *in vitro* studies showed that KT mutations at residues 25 and 26 (36) and Ala substitutions at any two or more MA-HBR basic residues (35) reduce RNA binding to MA. In cells, a reduction in tRNA populations bound to Gag relative to total RNA populations bound to Gag has been observed for KT mutations at residues 25 and 26 using photoactivatable ribonucleoside-enhanced crosslinking and immunoprecipitation (PAR-CLIP) followed by sequencing (PAR-CLIP-Seq) (47). However, the magnitude of the effect of the HBR mutations on the RNA binding ability of MA in cells remains to be determined. The second type of amino acid substitutions in MA-HBR that we introduced is Lys-to-Arg (KR) changes, which do not affect the overall basic charge of MA-HBR unlike the KT changes. Mutations that maintain the overall basic charge are expected to preserve the RNA binding ability, since liposome binding of a mutant Gag in which all basic residues of MA-HBR were switched (K->R and R->K) is still sensitive to RNA-mediated block of PC+PS liposome binding (52). However, this prediction has neither been tested in cells nor tested for specific HBR residues.

The comparison of HIV-1 Gag harboring the double amino acid substitutions in MA-HBR in this study revealed that indeed basic-to-neutral changes in two Lys residues are sufficient to diminish MA-RNA binding in cells. Furthermore, unlike the KT mutants, which did not bind RNA via MA in cells and were distributed to the PM as well as intracellular membranes, the KR mutants, which bound RNA, showed little or no binding to intracellular membranes regardless of their PM localization. Altogether, the strong correlation between MA-RNA binding and subcellular localization of Gag supports our working model that MA-RNA binding inhibits promiscuous localization of Gag, thereby ensuring Gag localization to the PM in the presence of PI(4,5)P2. The data obtained in this work also raise the possibility that the interaction of MA-HBR residues with PI(4,5)P2 modulates a step beyond targeting and binding of Gag to the PM.

## Results

### Gag derivatives with Lys-to-Arg (KR) changes in MA-HBR bind RNA more efficiently than those with Lys-to-Thr (KT) changes in cells

To compare the RNA binding capacity of MA, we introduced three modifications into Gag constructs. First is an MA amino acid substitution (1GA) that blocks N-terminal myristoylation and thereby prevents Gag from binding membranes. The previous cell-based PAR-CLIP study (47) was conducted using Gag constructs that can bind membranes, which would cause dissociation of RNA from MA according to the model described above (see Introduction). Therefore, we eliminated membrane binding so as to allow us to determine the total RNA binding capacity of Gag. Second, to focus on the RNA binding ability of the MA domain, we deleted most of the NC domain (delNC), which is the major RNA binding domain of Gag. Lastly, we fused YFP to the Gag C-terminus (Gag-YFP) to facilitate microscopy analysis of the same constructs analyzed for MA-RNA binding. The 1GA/delNC/Gag-YFP constructs containing KR or KT mutations in MA residues 25 and 26 (25/26KR and 25/26KT) or MA residues 29 and 31 (29/31KR and 29/31KT) (Fig 1) were then compared with isogenic constructs with the WT HBR sequence or a mutant HBR sequence in which all MA-HBR basic residues were switched to neutral residues (6A2T). To determine the amount of RNA bound to MA in cells, we employed a modified PAR-CLIP assay (47, 53) (Fig 2). HeLa cells were transfected with one of the constructs described above or a non-Gag control, pUC19, cultured in a medium containing 4-thiouridine (4-SU), a photoactivatable ribonucleoside analogue, and subsequently exposed to UV light, which crosslinks RNA-binding proteins and 4-SU-containing RNA bound to the proteins. Following cell lysis, Gag constructs were immunoprecipitated using HIV-Ig, and the RNA bound to the constructs was end-labelled with ^32^P prior to SDS-PAGE and electrotransfer to PVDF membrane. The RNA binding efficiency of the constructs was determined through comparison of the signal intensity of RNA by phosphorimager analysis versus the signal intensity of Gag constructs detected by immunoblotting on the same membrane. Consistent with previous results (46), the Gag construct bearing 6A2T MA did not bind RNA efficiently, showing ∼4-fold reduction in the amount of RNA bound relative to WT. Both the constructs bearing 25/26KT and 29/31KT MA bound significantly less RNA than the WT construct, similar to what was observed for 6A2T, indicating that both Lys 25 and 26 and Lys 29 and 31 contribute to MA-RNA binding to similar extent in cells. On the other hand, there was no statistically significant difference in the RNA binding efficiency between the 25/26KR MA and the WT MA. The Gag construct with the other KR changes, 29/31KR, also bound significantly more RNA compared to its KT counterpart, albeit less efficiently than WT. These results indicate that the RNA binding efficiency of MA is dependent on the overall positive charge of the MA-HBR basic patch. Additionally, the identity of basic amino acids at residues 29 and/or 31 also plays a role in WT-level RNA binding. Having observed differential effects on MA-RNA binding between KR and KT substitutions, we investigated below the correlation (or lack thereof) between MA-RNA binding and subcellular localization of Gag, first focusing on the effects of the substitutions at MA residues 25/26, and next on those at the residues 29/31.

**Fig 1.**
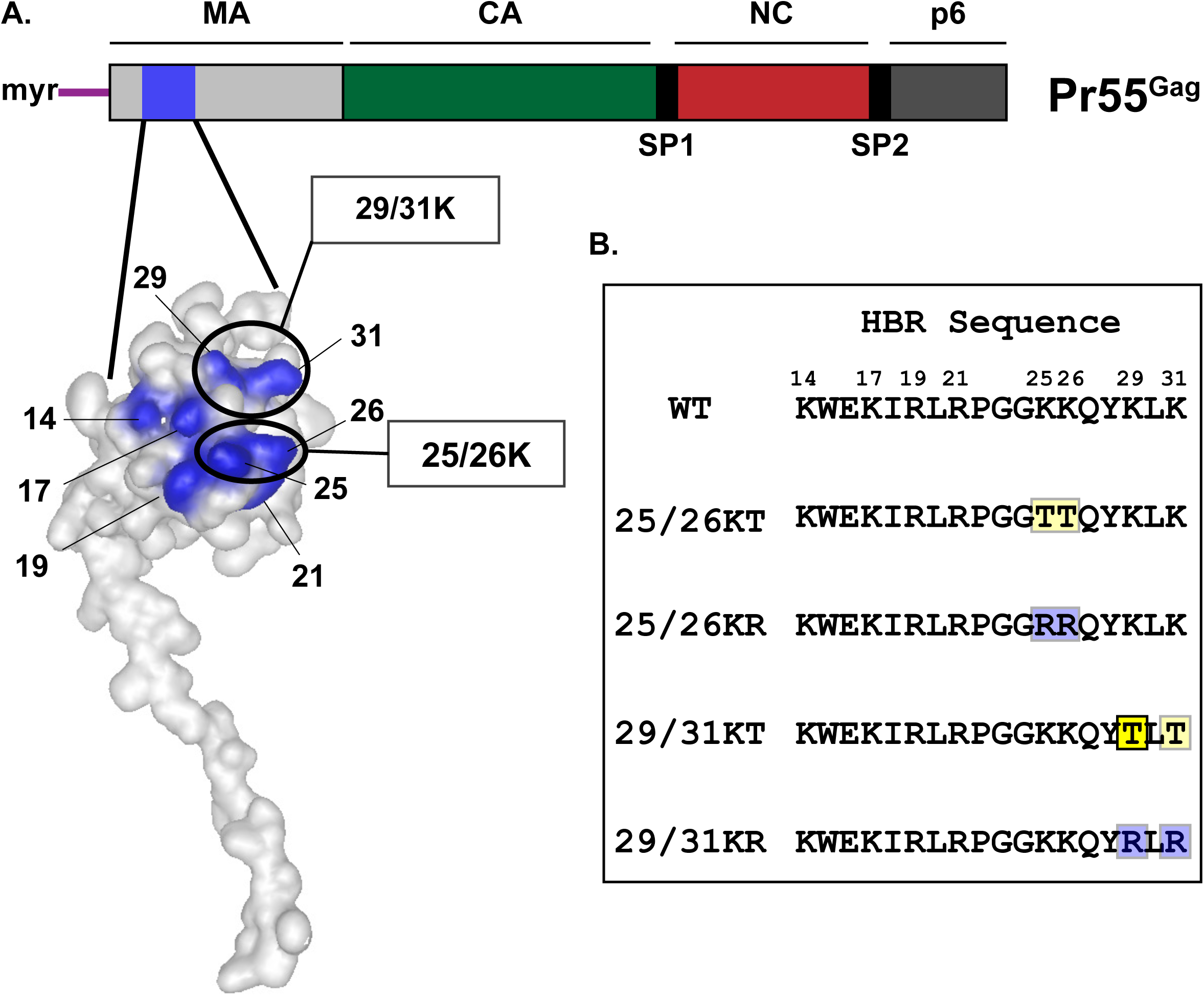
Amino acid substitutions introduced in MA-HBR. A: Schematic illustration of HIV-1 Gag with the structure of HIV-1 MA (PDB 2HMX) showing the basic residues of HBR in blue with residues mutated in this study circled. B: Sequences of HIV-1 Gag MA-HBR analyzed in this study are shown. Lys-to-Arg (KR) or Lys-to-Thr (KT) changes were introduced at MA residues 25 and 26 or 29 and 31.

**Fig 2.**
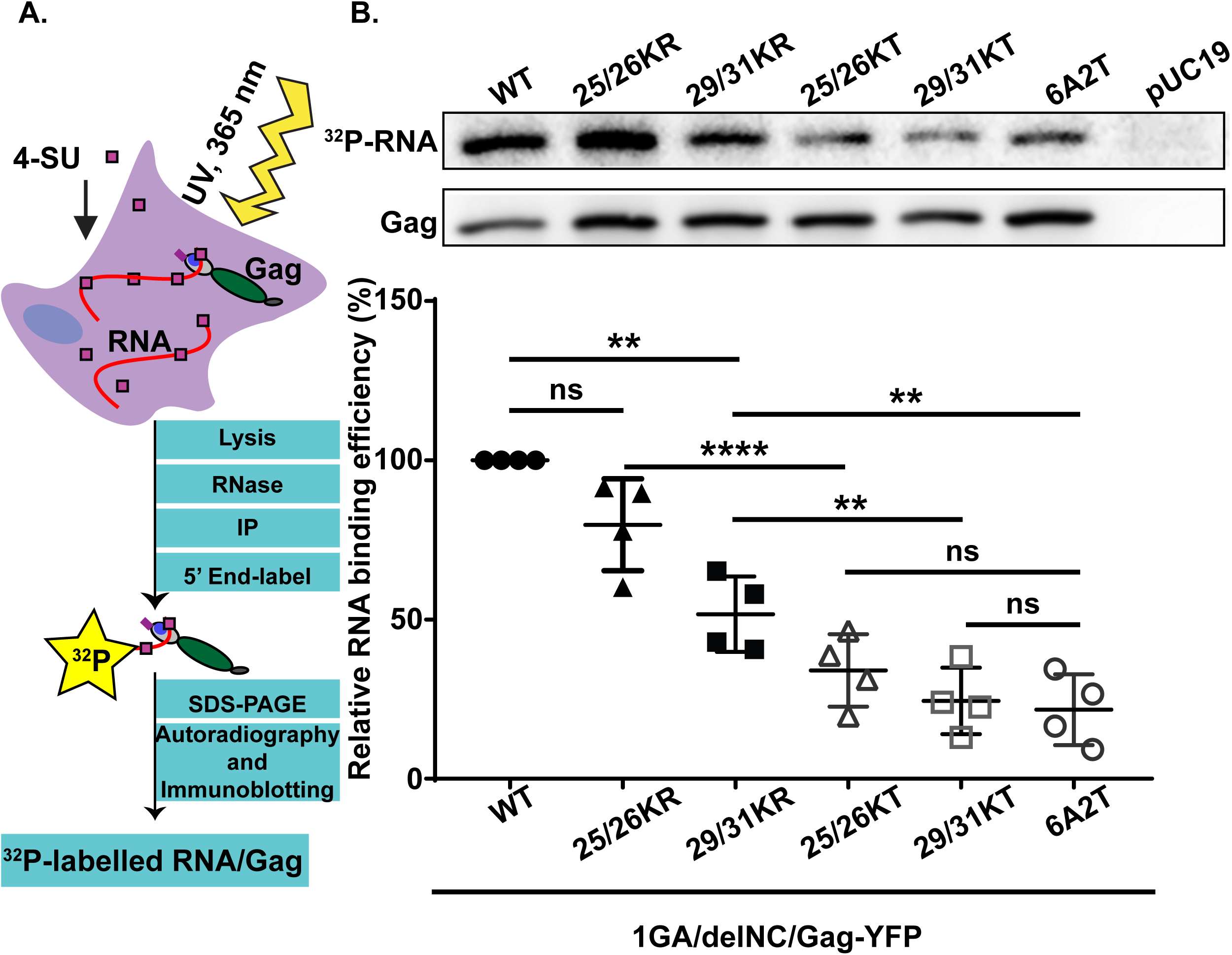
Gag derivatives with KR mutations in MA-HBR bind RNA efficiently compared to those with KT mutations. A: Schematic illustration of the PARCLIP assay. A CMV promoter-driven Gag-YFP construct lacking the myristoylation site (1GA) and most of the NC domain (delNC) was used as the backbone for the Gag derivatives analyzed in this assay. A mutant Gag with all MA-HBR basic residues switched to neutral Ala or Thr, 6A2T, was used as a negative control. To identify bands representing Gag, non-Gag plasmid, pUC19, was used as an additional control. The 1GA/delNC/Gag-YFP constructs were expressed in HeLa cells and crosslinked to 4-SU containing RNA in the cells. Gag proteins were recovered by immunoprecipitation, Gag-bound RNA was end-labeled with 32P, and signals for Gag proteins and Gag-bound RNA were detected by SDS-PAGE followed by immunoblotting and autoradiography, respectively. B: Representative results for 32P-labelled Gag-bound RNA detected by autoradiography and total Gag detected by immunoblotting and chemiluminescence are shown on the top. Relative RNA binding efficiency (%) of MA-HBR mutants was determined by quantifying the intensity of RNA signals normalized by the Gag band intensity on the same PVDF membrane. Results from 4 independent experiments are shown as means +/- standard deviations. P values were determined using Student’s t-test, using raw data. ns, not significant; **, P ≤ 0.01; ****, P ≤ 0.0001.

### Gag derivatives containing 25/26KR substitutions exhibit PM-specific localization unlike those with 25/26KT changes

To compare subcellular distribution of Lys 25/26 mutants, we expressed myristoylated and YFP-tagged Gag constructs bearing these changes, both in the full-length and delNC contexts, in HeLa cells and observed the YFP localization in cells (Fig 3). Based on the YFP distribution patterns, we identified 4 different phenotypes: (i) localization predominantly at the PM (green bar), (ii) localization to both PM and intracellular compartments (blue bar), (iii) localization to only intracellular compartments (orange bar), and (iv) only hazy cytosolic localization with no punctate intensities (pink bar). These 4 distribution patterns were also validated by comparison of the YFP signal with that of Alexa Fluor 594-conjugated Concanavalin A (ConA), which serves as the PM marker. As seen previously, the 25/26KT mutant bound both PM and intracellular membranes. In contrast, the 25/26KR mutant presented PM-specific localization similar to WT. Importantly for correlating the localization data with MA-RNA binding results (Fig 2), the subcellular localization patterns of delNC/Gag-YFP was similar to that of the corresponding full-length Gag-YFP for both WT and the MA-HBR mutants. When we examined the VLP release efficiency of full-length Gag-YFP construct, we found that despite the difference in subcellular localization, both the Lys 25/26 mutants displayed robust VLP release efficiency compared to WT (Fig 4). A previous study showed that 25/26KT Gag produced *in vitro* using rabbit reticulocyte lysates binds neutral PC-only liposomes efficiently, presumably due to enhanced myristate-driven hydrophobic interactions (32). Using the same liposome binding assay, we found that the 25/26KR mutant also exhibits increased hydrophobic interactions (Fig 5A, 5B). These data suggest that the enhanced VLP release efficiency of Gag constructs with the Lys 25/26 substitutions is likely a result of increased hydrophobic interactions (see Discussion).

**Fig 3.**
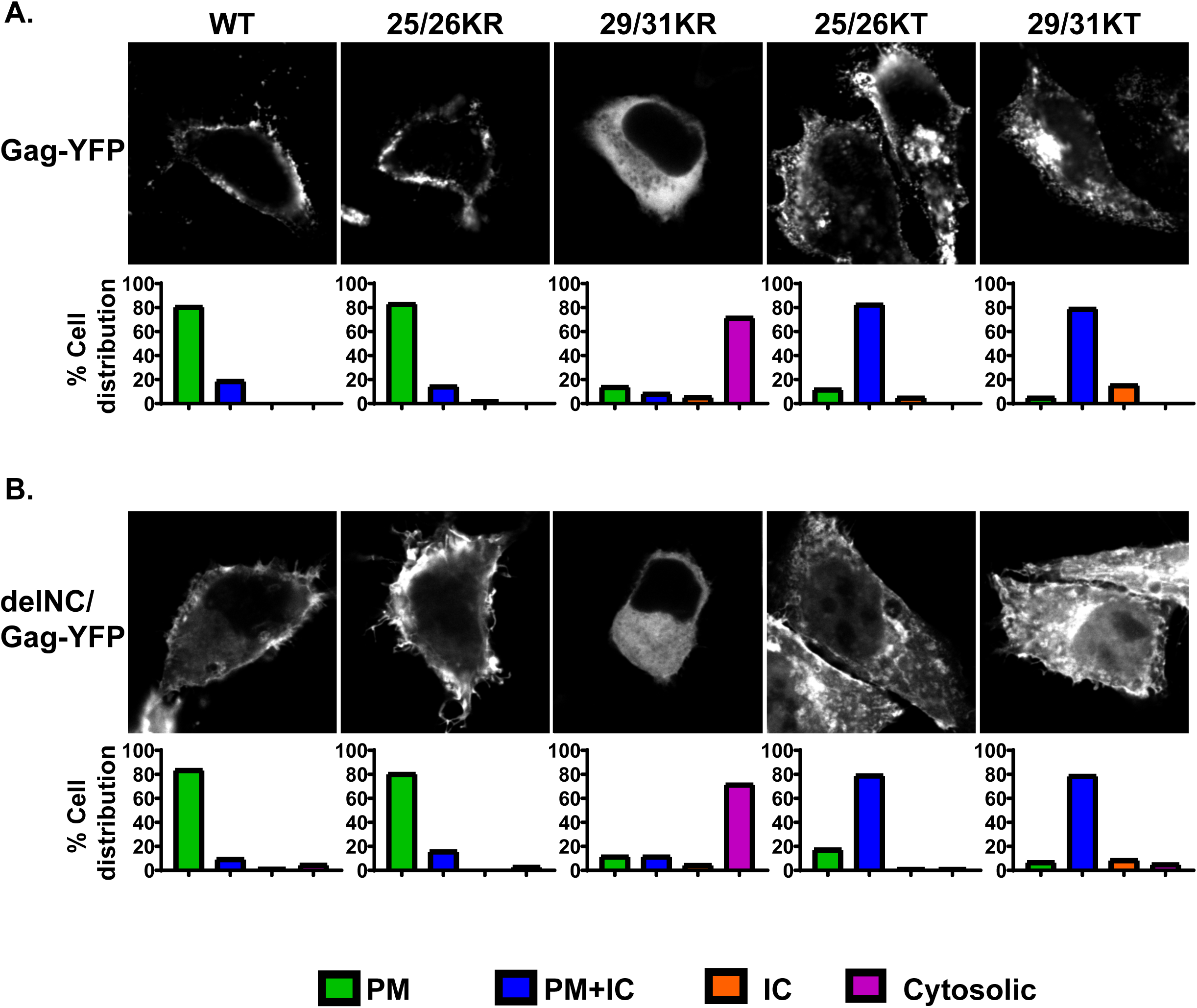
Gag derivatives containing KR substitutions in the MA-HBR region do not show promiscuous localization in cells unlike those with KT changes. HeLa cells were transfected with full-length Gag-YFP (A) or delNC/Gag-YFP (B), which contain WT MA sequence or 25/26KR, 25/26KT, 29/31KR, 29/31KT or 6A2T substitutions. At 14 hours post transfection, cells were stained with Alexa Fluor 594-conjugated concanavalin A (ConA), fixed with 4% paraformaldehyde in PBS, and analyzed using a fluorescence microscope. Note that subcellular distributions of delNC/Gag-YFP constructs mirror those of the corresponding full-length Gag-YFP constructs for both WT and MA-HBR mutants. Forty-two to 85 cells were analyzed per condition across 3 independent experiments. The localization patterns determined by epifluorescence microscopy were confirmed by confocal microscopy. Representative confocal images are shown in the top panels. Bar graphs in the bottom panels represent percentages of cell populations showing indicated patterns of subcellular distribution for each Gag-YFP construct. ConA staining was used as PM marker. PM, Plasma membrane; PM+IC, Plasma membrane + Intracellular compartments; IC, Intracellular compartments.

**Fig 4.**
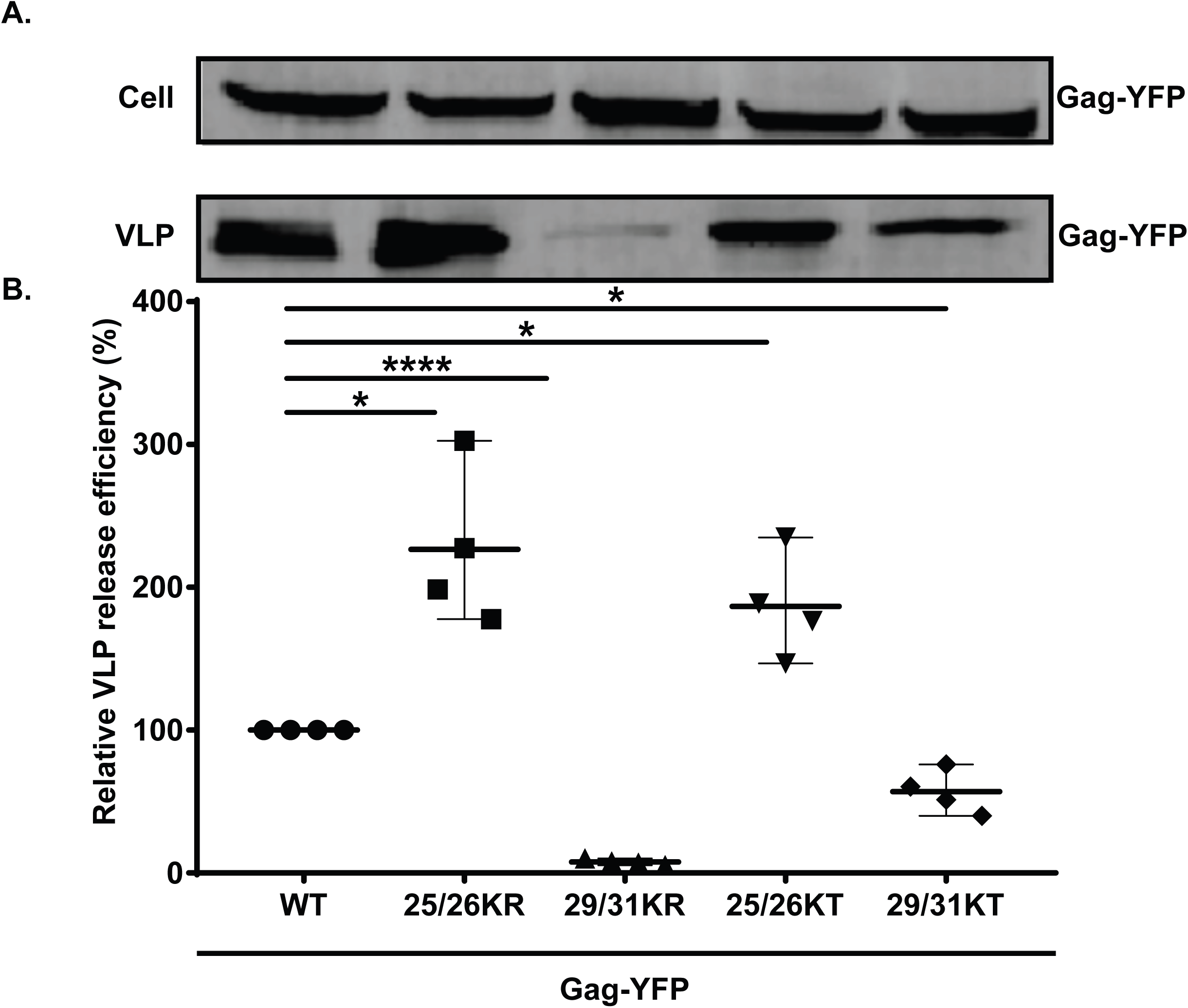
MA-HBR mutations alter VLP release efficiency. A: HeLa cells were transfected with WT Gag-YFP or Gag-YFP containing 25/26KR, 25/26KT, 29/31KR or 29/31KT changes. At 14 hours post transfection, cell and virus like particle (VLP) lysates were collected and subjected to SDS-PAGE, and Gag proteins were detected by immunoblotting using HIV immunoglobulin. B: Relative VLP release efficiency represents the amount of VLP-associated Gag as a fraction of total Gag present in VLP and cell lysates and is normalized to the VLP release efficiency of WT Gag. Results from 4 independent experiments are shown as means +/- standard deviations. The average VLP release efficiency of WT Gag was 8.3%. P values were determined from raw data using Student’s t- test. *, P ≤ 0.05; ****, P ≤ 0.0001.

**Fig 5.**
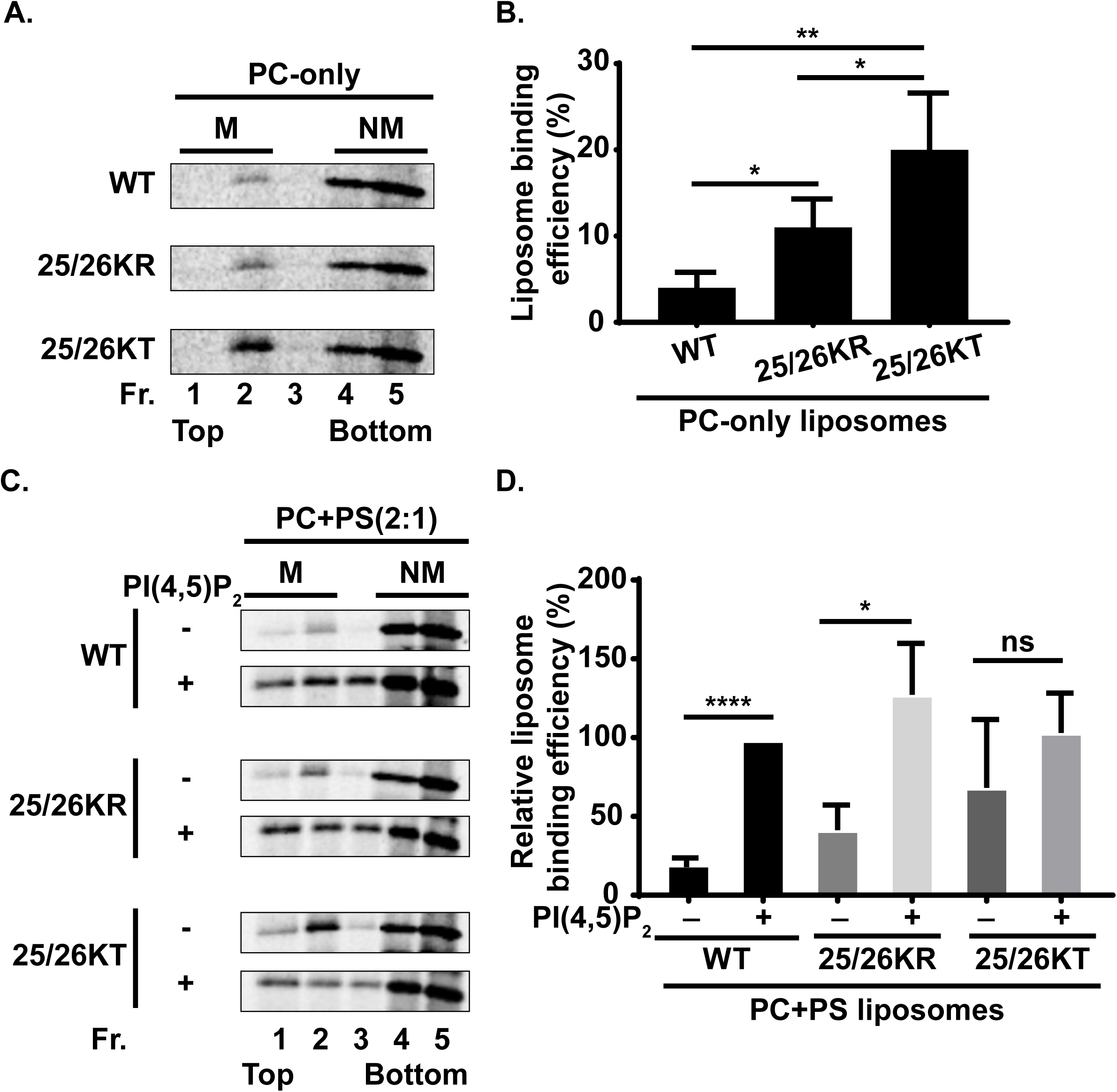
Liposomes binding efficiency of Gag derivatives containing Lys 25/26 substitutions. A: 35S-labeled HIV-1 WT Gag, 25/26KR Gag and 25/26KT Gag were synthesized in vitro using rabbit reticulocyte lysates and incubated with PC-only liposomes. The reaction mixtures were subjected to membrane flotation centrifugation, and a total of five 1-ml fractions were collected from each sample. M, membrane bound Gag; NM, non-membrane-bound Gag. B: The liposome binding efficiency was calculated as the percentage of membrane bound Gag (M) to the total Gag (M+NM) synthesized in the reaction. Results from four independent experiments are shown as means ± standard deviations. P values were determined by Student’s t test. *, P ≤ 0.05; **, P ≤ 0.01. C: 35S-labeled HIV-1 WT Gag, 25/26KR Gag and 25/26KT Gag were synthesized in vitro using rabbit reticulocyte lysates and incubated with liposomes [PC+PS (2:1)] or liposomes containing 7.25 mol% PI(4,5)P2 [PC+PS (2:1) + PI(4,5)P2]. The reaction mixtures were analyzed as in (A). M, membrane bound Gag; NM, non-membrane-bound Gag. D: The relative liposome binding efficiency was calculated as the percentage of membrane-bound Gag (M) to the total Gag (M+NM) synthesized in the reaction and normalized to the efficiency of WT Gag binding to PI(4,5)P2-containing liposomes. The average efficiency of WT Gag binding to PI(4,5)P2-containing liposome was 42.5%. Results from three independent experiments are presented as means ± standard deviations. P values were determined by Student’s t test. ns, not significant; *, P ≤0.05; ****, P ≤ 0.0001.

### 25/26KR Gag is dependent on PI(4,5)P2 for efficient membrane binding unlike 25/26KT Gag

While 25/26KR MA shows WT-level RNA binding and mediates PM-specific localization of Gag-YFP, unlike WT MA it engaged in increased hydrophobic interactions with membranes similar to its corresponding KT mutant MA (32) (Fig 5A, 5B). An earlier study has demonstrated that the 25/26KT mutant exhibits increased PI(4,5)P2-independent liposome binding (32). Therefore, we examined whether 25/26KR MA still shows PI(4,5)P2-dependent liposome binding. We found that the presence of PI(4,5)P2 significantly increased the liposome binding efficiency of the 25/26KR mutant but does not affect the 25/26KT mutant (Fig 5C, 5D). These results indicate that the 25/26KR mutant is dependent on PI(4,5)P2 for efficient membrane binding unlike the 25/26KT mutant even though both showed increased hydrophobic interactions relative to WT.

### Upon depletion of cellular PI(4,5)P2, 25/26KR Gag-YFP exhibits promiscuous subcellular localization

PM localization of WT Gag is dependent on PI(4,5)P2 (15, 26, 27). To investigate whether the PM localization of 25/26KR Gag is also dependent on PI(4,5)P2, we next examined the effect of expression of 5-phosphatase IV (5ptaseIV), which depletes cellular PI(4,5)P2, on subcellular distribution of Gag-YFP derivatives. We co-expressed WT Gag-YFP or Gag-YFP with Lys 25/26 substitutions with full-length 5ptaseIV (FL) or its catalytically inactive derivative (Δ1) (Fig 6). As previously reported (26), WT Gag-YFP lost PM-specific localization and displayed predominantly hazy cytosolic but sometimes intracellular punctate distribution when co-expressed with FL 5ptaseIV. The 25/26KT mutant showed predominantly promiscuous localization regardless of the presence of FL or Δ1 5ptaseIV, demonstrating that this MA substitution mutant is not dependent on PI(4,5)P2 for cellular membrane binding. In contrast, the localization of the 25/26KR mutant changed from predominantly PM-specific localization to more promiscuous localization when it was co-expressed with the FL but not Δ1 5ptaseIV. These results indicate that the PM-specific localization of the 25/26KR Gag is dependent on the presence of PI(4,5)P2 in the cell. Overall, our results suggest that in the case of the Lys 25/26 mutants, Gag that binds RNA efficiently maintains PI(4,5)P2-dependent PM-specific localization, while Gag that binds RNA poorly shows PI(4,5)P2-independent promiscuous localization. Thus, for Lys 25/26 substitutions, there is a correlation between RNA-binding and PI(4,5)P2-dependent PM-specific localization as observed with Gag chimeras containing heterologous retroviral MA domains (29).

**Fig 6.**
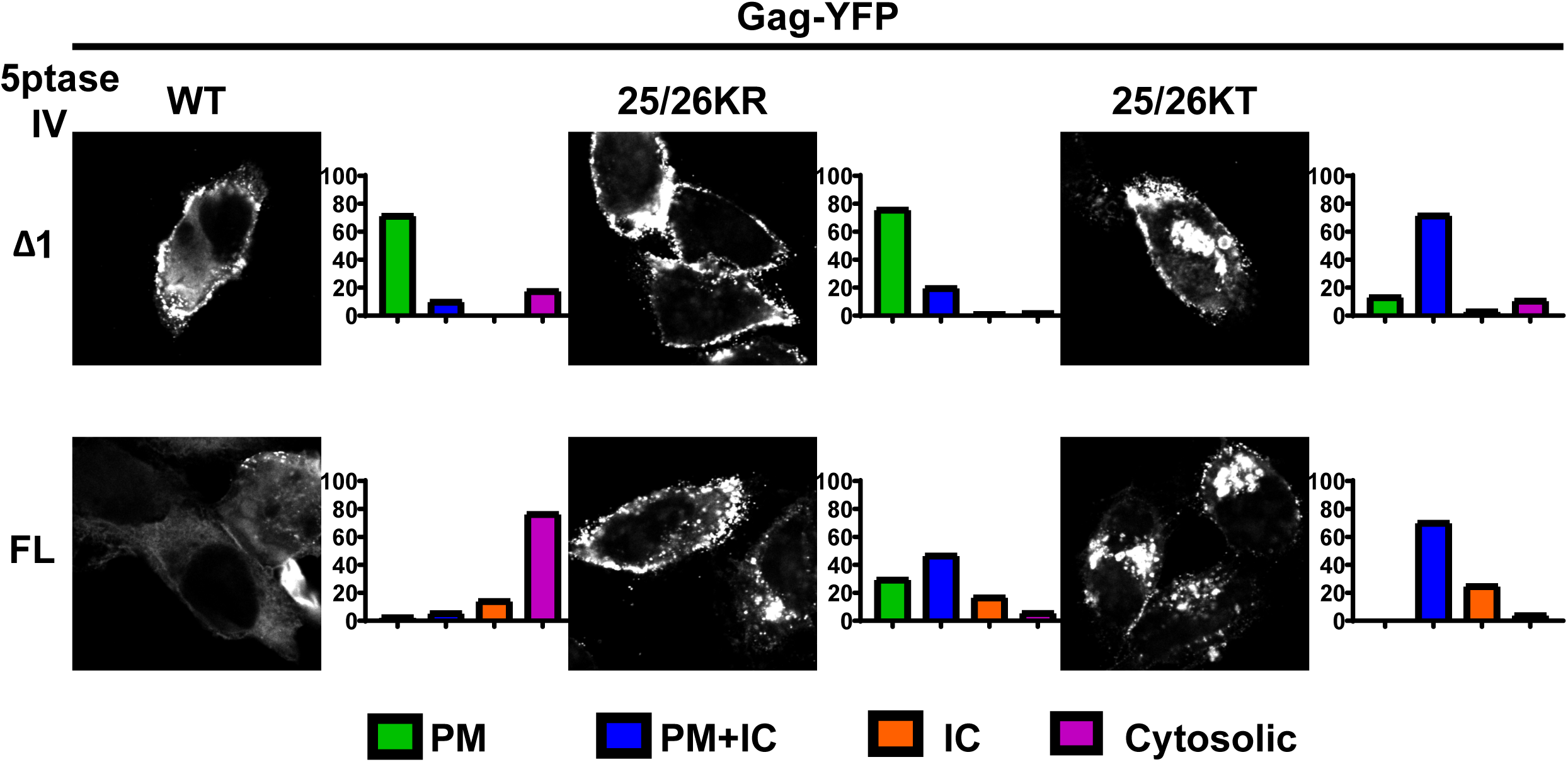
Depletion of cellular PI(4,5)P2 leads to promiscuous subcellular localization of 25/26KR Gag-YFP. HeLa cells were transfected with WT Gag-YFP, 25/26KR Gag-YFP or 25/26KT Gag-YFP along with myc-tagged 5ptaseIV FL or the catalytically inactive Δ1 derivative. At 14 hours post transfection, cells were stained with Alexa Fluor 594-conjugated ConA (not shown), fixed with 4% paraformaldehyde in PBS, permeabilized, immunostained with mouse monoclonal anti-Myc antibody and anti-mouse IgG conjugated with Alexa Fluor 405 (not shown), and analyzed using a fluorescence microscope. Only cells positive for both myc-tagged 5ptaseIV and Gag-YFP were included in the analysis. Sixty-nine to 134 cells were analyzed per condition across 3 independent experiments. Representative confocal images of Gag-YFP are shown. The localization patterns were examined as in Fig 3, and the percentages of cells showing indicated subcellular distribution patterns under each condition are shown in bar graphs. PM, Plasma membrane; PM+IC, Plasma membrane + Intracellular; IC, Intracellular.

### Gag derivatives containing 29/31KR substitutions show predominantly cytosolic distribution

Since 29/31KR MA exhibits significantly higher RNA binding than 29/31KT MA (Fig 2), we next sought to test whether RNA binding efficiency correlates with subcellular localization in the case of Gag constructs containing changes at MA residues 29/31 as were observed for Gag mutants with changes at residues 25/26. To this end, we first examined distribution of myristoylated full-length Gag-YFP and delNC/Gag-YFP derivatives (Fig 3). Consistent with previous reports, Gag-YFP with the 29/31KT changes displayed promiscuous localization with its signals present at both PM and intracellular membranes. Interestingly, the 29/31KR mutant displayed a predominantly cytosolic distribution of Gag-YFP indicating a loss of efficient membrane binding. It is important to note that unlike 29/31KT Gag-YFP, 29/31KR Gag-YFP showed negligible intracellular membrane binding despite having a higher positive charge than 29/31KT MA, which could mediate electrostatic interactions with acidic lipids more efficiently. We found that both delNC and full-length Gag-YFP showed analogous subcellular patterns for each Lys 29/31 mutant. These results, together with the results of the comparison between two Lys 25/26 mutants, strongly suggest that MA-RNA binding prevents Gag binding to intracellular membranes regardless of whether Gag can bind the PM. In congruence with the microscopy results, we found that the VLP release efficiency determined in the context of full-length Gag-YFP was significantly impaired for the 29/31KT mutant and almost abolished for the 29/31KR mutant (Fig 4).

### Replacing the NC domain with a heterologous dimerization motif enables 29/31KR Gag-YFP to localize specifically to the PM in a PI(4,5)P2-dependent fashion

Even though the lack of binding to intracellular membranes is a shared phenotype between 25/26KR Gag-YFP and 29/31KR Gag-YFP, 29/31KR Gag-YFP showed cytosolic distribution in most cells unlike 25/26KR Gag-YFP. We hypothesized that 29/31KR Gag, unlike WT and 25/26KR Gag, is unable to counteract the negative regulation of membrane binding by RNA that is bound to its MA-HBR. A previous study suggests that while overall positive charge of HBR is sufficient for RNA-mediated suppression of Gag-liposome binding irrespective of the identity of the residues, the identity of at least one of the basic residues is important for MA-PI(4,5)P2 interaction (52). In addition, a recent NMR study found that Lys 29 and 31 are important for binding of MA to PI(4,5)P2, even in an experimental system in which RNA is absent (49). Based on these observations, we reasoned that due to attenuated MA-PI(4,5)P2 binding, 29/31KR Gag is unable to alleviate RNA-mediated suppression of membrane binding. It is then conceivable, according to our model, that if membrane binding of this mutant Gag can be enhanced enough to offset the negative regulation imposed by MA-RNA interactions, then this mutant would localize specifically to the PM because MA-bound RNA would still prevent promiscuous membrane binding. Multimerization increases binding of HIV-1 MA to the PM as well as to liposomes of different compositions (48). Thus, we sought to test the hypothesis above by attempting to augment multimerization of 29/31KR Gag. Previous studies reported that a leucine zipper dimerization motif (LZ) promotes Gag multimerization more efficiently than the NC domain (55, 56). Therefore, in order to improve Gag multimerization, we replaced the NC domain of Gag with LZ (38, 57–60). We then expressed WT and the Lys 29/31 substitution mutants in the context of Gag-YFP and GagLZ-YFP in cells (Fig 7). We found that WT GagLZ-YFP predominantly localized at the PM as observed for WT Gag-YFP. The 29/31KT GagLZ-YFP showed a promiscuous localization like the 29/31KT Gag-YFP, indicating that merely replacing the NC domain with LZ did not lead to the plasma membrane-specific localization. Consistent with our prediction, the 29/31KR GagLZ-YFP demonstrated a markedly enhanced PM localization compared to the 29/31KR Gag-YFP. Importantly, this construct showed very little promiscuous localization unlike the 29/31KT GagLZ-YFP, suggesting that MA with 29/31KR changes retains the ability to ensure PM-specific localization. Unexpectedly, the 29/31KR GagLZ-YFP still failed to release VLPs efficiently (Fig 8), suggesting that MA residues 29 and 31 play an additional role in assembly beyond PM-specific binding. We then wanted to test directly in cells whether LZ-driven PM localization is still PI(4,5)P2-dependent. To this end, we co-expressed GagLZ-YFP constructs with FL and Δ1 5ptaseIV enzyme (Fig 9). We found that both the WT GagLZ-YFP and 29/31KR GagLZ-YFP lost the PM-specific localization when the FL 5ptaseIV enzyme was co-expressed. These results suggest that the WT GagLZ-YFP and the 29/31KR GagLZ-YFP bind the PM in a PI(4,5)P2-dependent manner. Interestingly, in the presence of FL 5ptaseIV, WT GagLZ-YFP showed promiscuous localization to intracellular compartments and the PM, whereas 29/31KR GagLZ-YFP showed predominantly cytosolic localization as observed for WT Gag-YFP (Fig 6). Overall, the presented data suggest that when MA-HBR in Gag binds RNA, Gag localizes specifically to the PM if it can counteract the RNA-mediated block on membrane binding either via interaction with PI(4,5)P2, which occurs natively in the case of WT and 25/26KR Gag, or when enhanced by improved multimerization as observed for 29/31KR GagLZ. In contrast, when Gag is deficient in MA-RNA binding, it fails to localize specifically to the PM and instead, binds promiscuously to different cellular membrane compartments. Altogether the analyses of Lys 25/26 and Lys 29/31 substitutions in MA-HBR revealed a strong correlation between MA-RNA binding and suppression of non-specific Gag localization to intracellular membranes.

**Fig 7.**
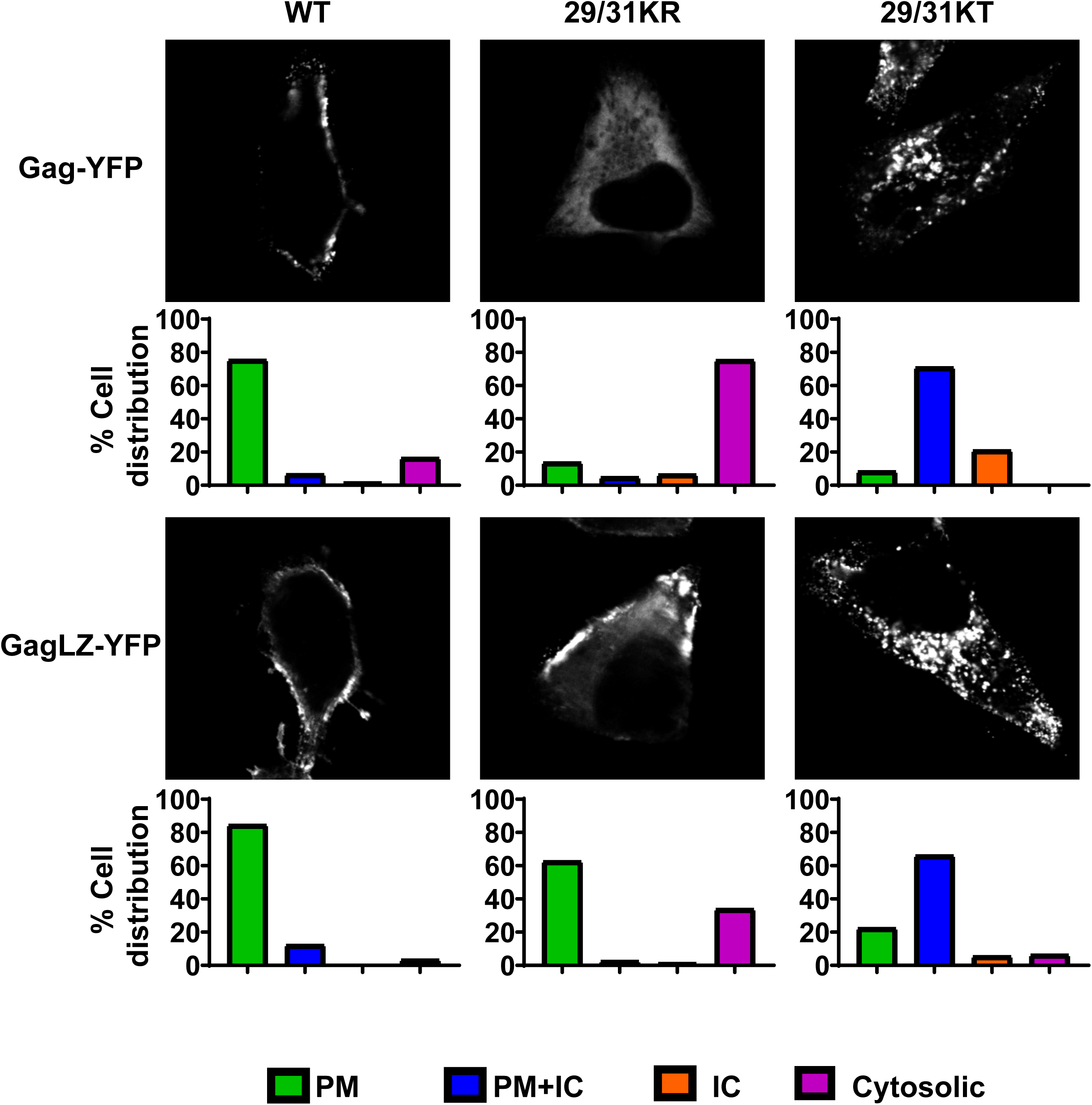
29/31KR Gag exhibits increased PM-specific localization when the NC domain is exchanged for leucine dimerization motif (LZ). HeLa cells were transfected with WT Gag-YFP, WT GagLZ-YFP or their derivatives with 29/31K substitutions. At 14 hours post transfection, cells were stained with Alexa Fluor 594-conjugated concavalin A (ConA, not shown), fixed with 4% paraformaldehyde in PBS, and analyzed using a fluorescence microscope. Sixty-one to 125 cells were analyzed per condition across 3 independent experiments. Representative confocal images of Gag-YFP are shown (top panels). The localization patterns were examined as in Fig 3, and the percentages of cells showing indicated subcellular distribution patterns are shown in bar graphs (bottom panels). PM, Plasma membrane; PM+IC, Plasma membrane + Intracellular; IC, Intracellular.

**Fig 8.**
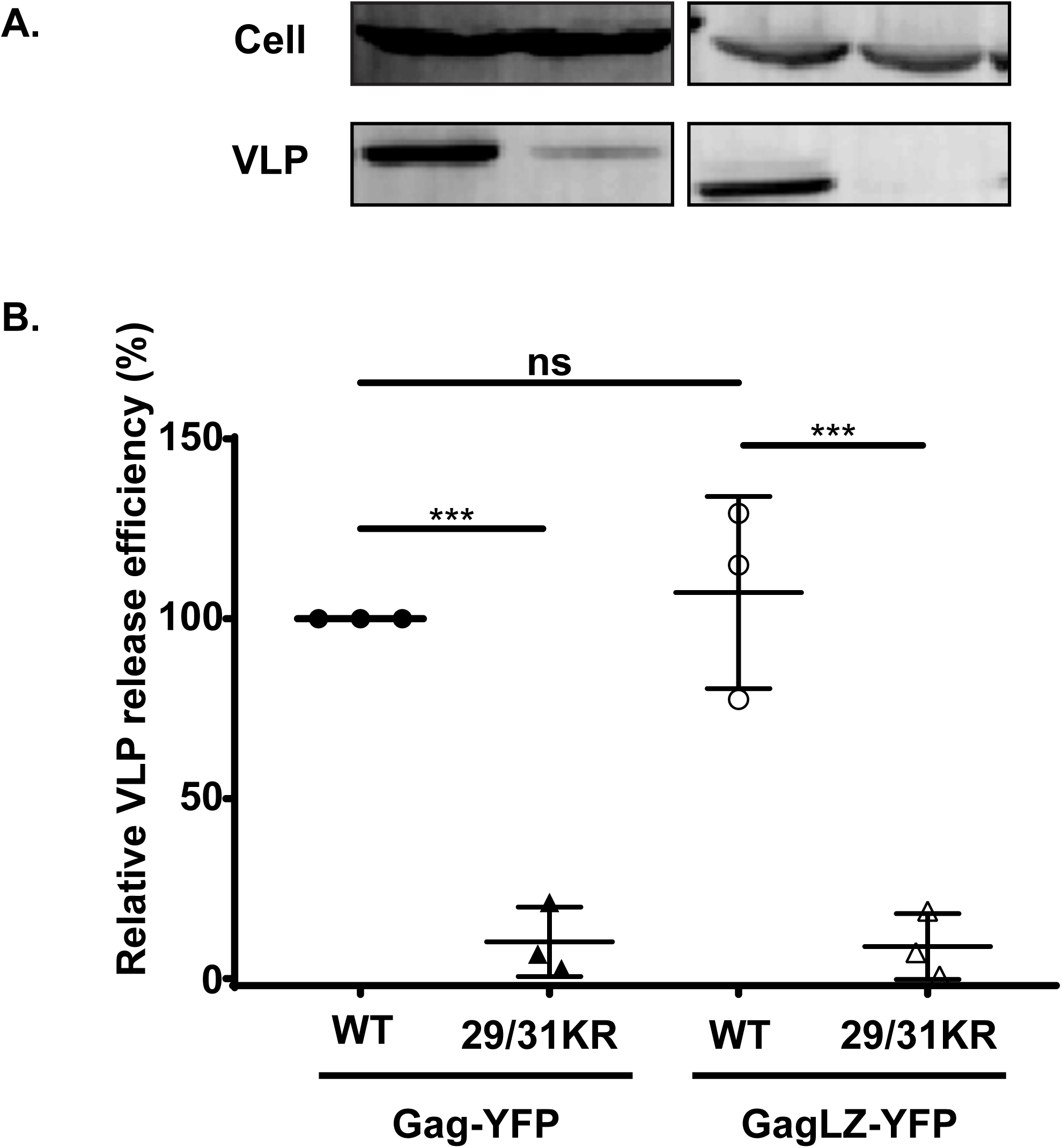
29/31KR GagLZ-YFP fails to release VLP. A: HeLa cells were transfected with Gag-YFP or GagLZ-YFP containing WT MA or 29/31KR MA sequences. At 16 hours post transfection, cell and VLP lysates were collected and analyzed as in Fig 4. B: Relative VLP release efficiency represents the amount of VLP-associated Gag as a fraction of total Gag present in VLP and cell lysates and is normalized to the VLP release efficiency of WT Gag. Results from 3 independent experiments are shown as means +/- standard deviations. The average VLP release efficiency of WT Gag was 9.6%. ns, not significant; ***, P ≤ 0.001.

**Fig 9.**
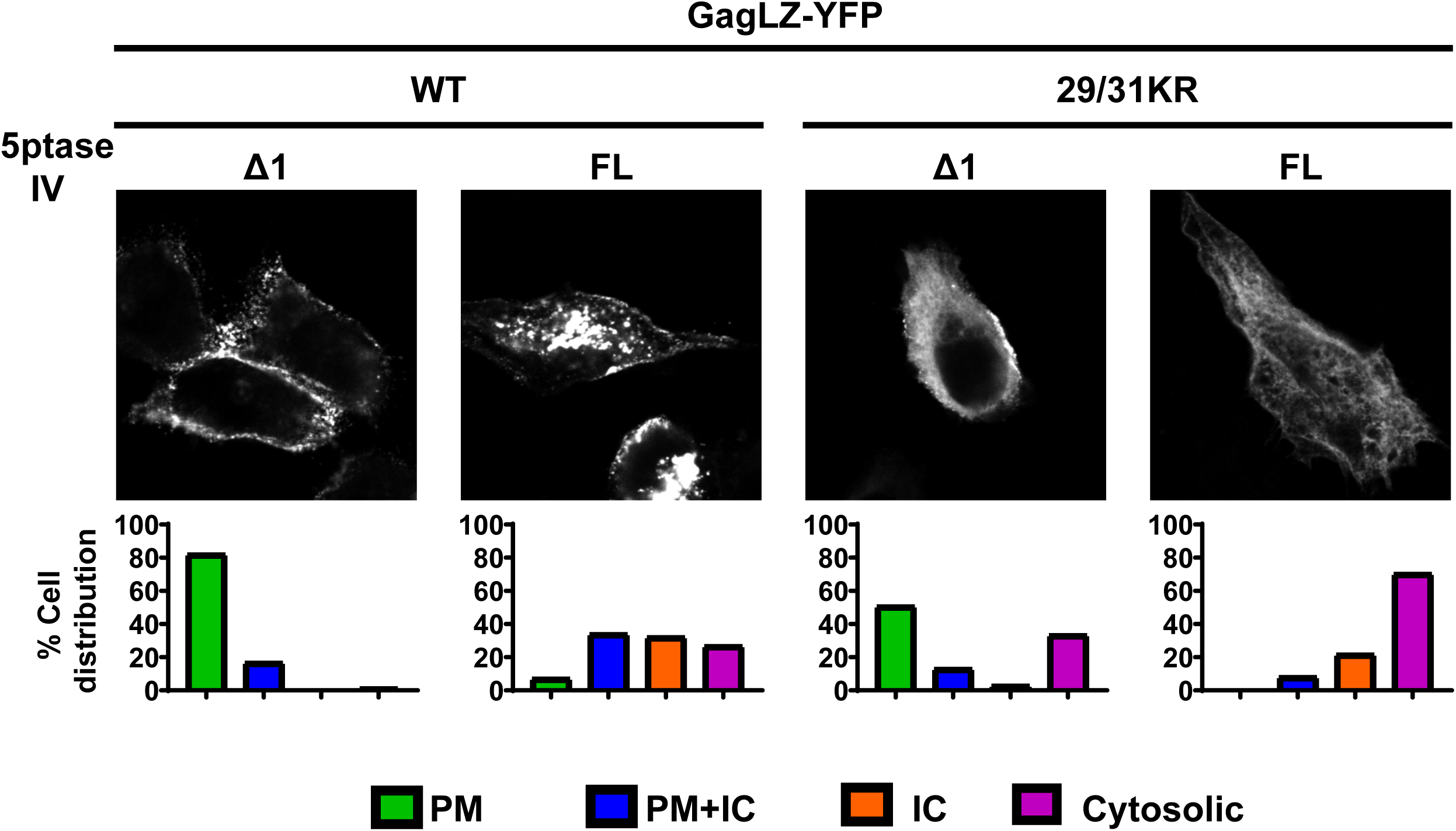
29/31KR GagLZ-YFP localizes to the PM in a PI(4,5)P2-dependent manner. HeLa cells were transfected with WT GagLZ-YFP or 29/31KR GagLZ-YFP along with myc-tagged 5ptaseIV FL or the catalytically inactive Δ1 derivative. At 14-16 hours post transfection, cells were stained with Alexa Fluor 594-conjugated ConA (not shown), fixed with 4% paraformaldehyde in PBS, immunostained with mouse monoclonal anti-Myc antibody and anti-mouse IgG conjugated with Alexa Fluor 405 (not shown), and analyzed as in Fig 6. Thirty-seven to 78 cells were analyzed per condition across 3 independent experiments. Representative confocal images of Gag-YFP are shown (top panels). The localization patterns were examined as in Fig 3, and the percentages of cells showing indicated subcellular distribution patterns are shown in bar graphs (bottom panels). PM, Plasma membrane; PM+IC, Plasma membrane + Intracellular; IC, Intracellular.

## Discussion

In the current study, we examined the effects of amino acid substitutions in MA-HBR on the relative amount of RNA bound to HIV-1 Gag in cells and on the subcellular localization of Gag. We found that MA-HBR mutants that displayed a promiscuous subcellular localization (i.e., 25/26KT Gag and 29/31KT Gag) were incapable of binding RNA efficiently, consistent with earlier observations obtained with Gag chimeras containing heterologous retroviral MA sequences (29). In contrast, MA-HBR mutants that bound RNA efficiently did not show a promiscuous localization; they either localized specifically to the PM (25/26KR Gag) or mainly remained in the cytosol (29/31KR Gag). The latter presented PM-specific and not promiscuous localization when its membrane binding ability was enhanced via LZ-mediated multimerization. The correlation established between MA-RNA binding and the lack of Gag localization to intracellular membranes in this study further provides support for our current working model of HIV-1 Gag assembly in which RNA-mediated inhibition of membrane binding of Gag prevents non-specific localization of Gag to membranes other than the PM.

The liposome binding experiments using neutral liposomes indicate that 25/26KR Gag engages in greater hydrophobic interactions with membranes than WT Gag, likely due to increased myristoyl exposure relative to WT Gag. Combined with analogous results for 25/26KT Gag, first observed in our previous study (32), these results support that Lys 25 and 26 play a role in myr exposure regulation. Despite the enhanced hydrophobic interactions with membranes, 25/26KR Gag maintains PM-specific localization like WT Gag. However, the outcomes of PI(4,5)P2 depletion differ between WT and 25/26KR Gag. Upon FL 5ptaseIV expression, WT Gag loses membrane binding and exhibits predominantly cytosolic distribution, whereas 25/26KR Gag retains membrane binding albeit localizing promiscuously in the majority of the Gag-YFP expressing cells. Therefore, RNA binding alone is likely sufficient for preventing promiscuous binding of WT Gag to intracellular membranes, but the presence of PI(4,5)P2 is necessary when membrane binding is enhanced beyond the WT level. The two different outcomes of PI(4,5)P2 depletion also highlights the two functions of PI(4,5)P2, enhancing membrane binding of Gag and supporting PM-specific localization.

We found that 25/26KR Gag exhibits increased VLP release efficiency compared to WT Gag, likely due to enhanced membrane binding as discussed above. Therefore, the Lys-to-Arg mutation at residues 25 and 26 should confer fitness advantage to HIV-1. However, Lys 25 and 26 are highly conserved across various clades of HIV-1 (61). We speculate that Arg, instead of Lys, at positions 25 and 26 may negatively affect other aspects of the HIV-1 life cycle. Similar to 25/26KR Gag, 25/26KT Gag also shows superior VLP release efficiency even though 25/26KT Gag does not localizes specifically to the PM unlike its KR counterpart. The high VLP release efficiency of 25/26KT Gag may be due to even more robust hydrophobic interactions with membranes than 25/26KR Gag (Fig 5), which may make up for the Gag mislocalization. The increased hydrophobic interactions of 25/26KT Gag, which does not bind RNA, is consistent with the observation that removal of RNA increases hydrophobic interactions in a myr-dependent manner (32).

We found that RNA binding of 29/31KR Gag was not as efficient as WT Gag although it was better than that of 29/31KT Gag. While we observed previously that RNA sensitivity of Gag-liposome binding is maintained when all Lys and Arg in MA-HBR are switched with each other (52), the presence of Lys, but not Arg, at residues 29 and 31 seems to be important for WT-level RNA binding of Gag. While both Arg and Lys are positively charged basic amino acids, Arg has a terminal guanidium group, while Lys has a single terminal amino group. These groups also differ in their geometry (i.e., planar versus tetrahedral). Therefore, it is possible that Lys and Arg participate in different number and/or orientation of electrostatic interactions with RNA or lipids (62–65).

Even though 29/31KR Gag showed reduced RNA binding, this level of MA-RNA binding was sufficient for suppressing binding to intracellular membranes. It is noteworthy that 29/31KR Gag failed to bind even the PM in most cells. Strongly cationic model proteins have been shown to associate with the PM (54). However, in the case of HIV-1 Gag, the high positive charge retained in 29/31KR MA is insufficient for the PM localization, likely due to the RNA-mediated block. As mentioned above, Lys 29 and 31, especially 31, are important for PI(4,5)P2 interaction even in the absence of RNA (49). Based on this knowledge, we speculate that due to inefficient interaction with PI(4,5)P2, 29/31KR Gag is unable to overcome the negative regulation by RNA. Therefore, the balance between MA-RNA and MA-PI(4,5)P2 binding, which is carefully orchestrated for WT Gag, is tipped in the favor of MA-RNA binding in case of 29/31KR Gag. It is likely that the enhanced multimerization of 29/31KR Gag through the LZ dimerization motif restores the balance toward MA-PI(4,5)P2 binding, leading to PM-specific localization of 29/31KR GagLZ. Probable mechanisms through which LZ may promote membrane binding may be an avidity effect, induction of myristate exposure (22), a reduction in the amount of RNA bound to MA associated with myristate exposure (66), or some combination of these factors.

Notably, there is no commensurate increase in VLP release for 29/31KR GagLZ compared to 29/31KR despite the dramatic shift in Gag localization from the cytosol to the PM. It is conceivable that the 29/31KR mutation causes not only a membrane-binding defect but an additional cryptic defect that is revealed only in the LZ context. Since 29/31KR GagLZ is recruited to the PM, the defect is likely at an assembly step after targeting and binding of Gag to the PM. Analogous to our observation, a recent study found that when the MA domain of HIV-1 Gag was replaced with the N-terminal PI(4,5)P2 binding region of avian sarcoma virus Gag, the chimeric Gag localized to the PM but did not release VLP efficiently (67). The order of Gag-PM binding and multimerization into Gag puncta varies between different retroviruses (68). The majority of HIV-1 Gag is directly recruited to Gag puncta at the PM from the cytosol rather than after binding elsewhere on the PM (68, 69). Thus, it is possible that a change in this order due to the LZ substitution of NC may have made a step in VLP assembly, such as membrane curvature, more dependent on the native sequence of MA-HBR. It is known that unsaturated acyl chains favor membrane curvature (70). Previous *in vitro* studies have shown that HIV-1 Gag is sensitive to acyl chain saturation of phospholipids and prefers PI(4,5)P2 with unsaturated acyl chains (71, 72). Acute PI(4,5)P2 depletion studies revealed that Gag continues to interact with PI(4,5)P2 after initial membrane binding of Gag and Gag multimerization (73). Altogether, we speculate that the orientation of MA basic amino acids at the interface between the PM and Gag lattice may play a key role in interactions with PI(4,5)P2 or other acidic lipids with different acyl chains and hence, in promotion of membrane curvature.

Overall, this paper provides a strong correlation between the ability of HIV-1 Gag to bind RNA via MA and the specificity of Gag localization to the PM. In the absence of PI(4,5)P2, WT Gag and 29/31KR GagLZ remain in the cytosol (Figs 6 and 9). Therefore, RNA binding is likely sufficient for preventing non-specific binding to other membranes for these Gag proteins. On the other hand, 25/26KR Gag and WT GagLZ, which are likely to have higher affinities to lipid bilayers due to increased myristoyl exposure or avidity, show promiscuous localization upon PI(4,5)P2 depletion (Fig 6 and 9). In these cases, both RNA and PI(4,5)P2 likely work in conjunction to prevent promiscuous localization. Altogether, the comparisons between the MA HBR mutants indicate the role of MA-bound RNA as an important player in targeting Gag specifically to the PM.

## Methods

### Plasmids

pCMVNLGagPolRRE Gag-YFP, which expresses HIV-1 Gag_NL4-3_ fused to YFP in an HIV-1 Rev-dependent manner, was constructed as described previously for pCMVNLGagPolRRE Gag-mRFP (74) using standard molecular cloning techniques. pCMVNLGagPolRRE delNC/Gag-YFP was generated by replacing the SpeI-XmaI region of pCMVNLGagPolRRE (75) with the corresponding region of pNL4-3/delNC/Gag-YFP described previously (76). 1GA mutation was introduced into the pCMVNLGagPolRRE delNC/Gag-YFP constructs by PCR mutagenesis as described before (74) pCMVNL GagLZ-YFP (previously named pCMVNLGag Venus LZ) was described in prior reports Gag (29, 52). pGEMNLNR, an expression vector used for *in vitro* transcription and translation of Gag, was described previously (26). To create pCMVNLGagPolRRE 25/26KR Gag-YFP and pCMVNLGagPolRRE 29/31KR Gag-YFP, the corresponding MA-HBR point mutations were introduced using PCR mutagenesis into pCMVNLGagPolRRE Gag-YFP. pCMVNLGagPolRRE 25/26KT Gag-YFP and pCMVNLGagPolRRE 29/31KT Gag-YFP were constructed by replacing the SpeI-BssHII region of pCMVNLGagPolRRE Gag-YFP with corresponding region of pGEMNLNR 25/26KT Gag and pGEMNLNR 29/31KT Gag (26, 32). pCMVNLGagPolRRE delNC/Gag-YFP containing the MA-HBR mutations, i.e., 25/26KR, 25/26KT, 29/31KR and 29/31KT, were constructed using standard molecular cloning techniques. pCMVNLGagPolRRE 6A2T delNC/Gag-YFP was created by replacing the SpeI-BssHII region of pCMVNLGagPolRRE delNC/Gag-YFP with corresponding region of pGEMNLNR 6A2T Gag (26, 32). pGEMNLNR containing the MA-HBR mutations, i.e., 25/26KR and 29/31KR, were constructed using standard molecular biology techniques. pCMVNL GagLZ-YFP containing the MA-HBR mutations, i.e., 25/26KR, 25/26KT, 29/31KR and 29/31KT; and pGEMNLNR containing the MA-HBR mutations, i.e., 25/26KR and 29/31KR, were constructed using standard molecular cloning techniques.

pCMV-Rev was described previously (74) (kindly provided by S. Venkatesan, National Institutes of Health). A mammalian expression plasmid encoding myc-tagged 5-phosphatase IV (FL 5ptaseIV), pcDNA4TO/Myc5ptaseIV, and its catalytically inactive Δ1 derivative (Δ1 5ptaseIV) were previously described (26, 77). pCMV-Vphu, which encodes the codon-optimized version of the HIV-1 Vpu gene, was previously described (78) (a kind gift from K. Strebel).

### Cells and transfection

HeLa cells were cultured as described previously (32). For microscopy, 30,000 HeLa cells were plated into each well of eight-well chamber slides (Lab-Tek; Nalgene Nunc International), cultured for 24 h, and transfected with plasmids encoding indicated Gag-YFP derivatives using Lipofectamine 2000 (Invitrogen) as per the manufacturer’s instructions. For VLP release and PAR-CLIP experiments, 4 × 10^5^ cells were plated into each well of six-well plates (Corning), cultured for 24 h, and transfected as described above.

### VLP release assay and immunoblotting

VLP release assays were carried out as described previously (32). HIV-1 Gag in cell and VLP lysates was detected by immunoblotting using HIV immunoglobulin (HIV-Ig; AIDS Research and Reference Reagent Program) as a primary antibody. Alexa Fluor 488-conjugated goat anti-human IgG (Invitrogen) was used for the detection of primary antibodies for HIV-1 Gag. Fluorescence signals were detected and quantified using a Typhoon Trio imager (GE Healthcare).

### Immunostaining and fluorescence microscopy

Hela cells transfected with plasmids expressing Gag-YFP were incubated for 1 min at room temperature with Alexa Fluor 594-conjugated Concanavalin A (ConA; Invitrogen) for visualization of the plasma membrane cells 14-16 hours post transfection. Cells were then fixed with 4% paraformaldehyde (PFA) in PBS. The expression of 5ptaseIV (both full-length [FL] and Δ1) in cells was detected by immunostaining with mouse anti-myc antibody (9E10; Santa Cruz Biotechnologies) after permeabilizing the cells. For quantitative analysis of Gag localization phenotypes, images of about 20 fields were recorded using an Olympus IX70 inverted fluorescence microscope at 60x magnification, and a range of 37 to 134 cells were evaluated for Gag localization pattern. For 5ptaseIV coexpression studies, only cells that were positive for both Gag and 5ptaseIV (either FL or Δ1) were chosen for evaluation. Confocal microscopy using Leica SP5 Inverted confocal microscope was carried out to further verify localization patterns determined by epifluorescence microscopy.

### Liposome-binding assay

Preparation of liposomes, *in vitro* Gag translation, and sucrose gradient flotation centrifugation were performed as described previously (32). In brief, full-length myristoylated Gag was translated *in vitro* using Promega’s TnT Sp6 Coupled Reticulocyte Lysate System and incubated with sonicated liposomes composed of a neutral lipid, phosphatidylcholine, and an acidic lipid, phosphatidylserine, in a 2:1 molar ratio (PC+PS [2:1] liposomes) with or without addition of 7.25 mol% PI(4,5)P2. Following sucrose flotation centrifugation, five fractions were collected with the top two fractions representing the membrane bound, floating Gag. Following SDS-PAGE, Gag bands were detected by phosphorimager analysis and quantified using ImageQuant 1D TL v8.1 image analysis software. Liposome binding efficiency (%) was calculated as the percentage of membrane bound Gag versus the total Gag in all 5 fractions.

### PAR-CLIP assay

To determine the amount of cellular RNA bound to transfected Gag, the PAR-CLIP assay was performed as described before (47, 53) with some modifications. Briefly, in a 6-well plate, 4X10^5^ HeLa cells were cultured in DMEM supplemented with 5% fetal bovine serum in the presence of penicillin and streptomycin antibiotics. The cells were then transfected with either a plasmid encoding 1GA/delNC/Gag-YFP containing the indicated MA sequence or pUC19. The cells were cultured for 24 hrs post transfection before addition of fresh media containing 100 µM 4-thiouridine (4-SU, Sigma), a photo-crosslinkable ribonucleoside analogue. After 16 hours, the cells were exposed to 354nm UV light at 300mJ/cm^2^ to crosslink RNA and RNA-binding proteins. Cells were lysed with NP-40 lysis buffer (50mM HEPES buffer [pH 7.5], containing 150mM KCl, 2mM EDTA, 0.5% [v/v] NP40 substitute, 0.5mM DTT, and supplemented with protease inhibitor cocktail), and the lysate was treated with 20U/ml RNase A (Promega) and 60U/ml RQ1 DNase (Promega). Gag was immunoprecipitated by incubating the cell lysates with HIV-Ig bound to Dynabeads^TM^ Protein G (Invitrogen) for 1 hour on a rotating shaker at 4°C. The RNA bound to Gag was first dephosphorylated at the 5’ end with calf intestinal alkaline phosphatase (New England BioLabs, NEB) followed by T4 polynucleotide kinase-catalyzed (NEB) rephosphorylation of the RNA at the 5’end with ^32^P (ATP-[γ-32P], PerkinElmer). The Gag-RNA complex was eluted from Dynabeads by heating in PARCLIP elution buffer (50mM Tris HCL pH 6.8, containing 2mM EDTA-NaOH 10% [v/v] Glycerol, 2% [v/v] SDS, 100mM DTT, 0.1% [w/v] Bromophenol Blue) for 5 min at 95°C with shaking at 1400 RPM. After resolving the bands by SDS-PAGE and subsequent electrotransfer to a PVDF membrane, the membrane was probed with HIV-Ig as the primary antibody and corresponding secondary antibody conjugated to horseradish peroxidase (goat anti-human IgG, Invitrogen). Gag-specific bands were detected after incubation with SuperSignal West Pico chemiluminescent substrate (Pierce) on Syngene Pxi multiapplication imager, and the intensity of the bands were measured using GeneTools analysis software. The ^32^P end-labeled RNA bands were also detected on the same membrane using autoradiography on the Typhoon FLA 9500 laser scanner and quantified using ImageQuant 1D TL v8.1 image analysis software. The relative RNA binding efficiency (%) was calculated as the percentage of total RNA band intensity versus the total Gag band intensity for each construct and compared to the normalized value for WT 1GA/delNC/Gag-YFP.

### Statistical analysis

Student’s t-tests were performed using Graphpad Prism. *P* values of <0.05 were considered statistically significant.

## ACKNOWLEDGMENTS

We thank our fellow laboratory members for helpful discussions and critical reviews of the manuscript. We thank S. Kutluay for her helpful comments in the optimization of the PAR-CLIP technique. The following reagent was obtained through the AIDS Research and Reference Reagent Program, Division of AIDS, NIAID, NIH: HIV Ig (from NABI and NHLBI). The microscopy data presented in this study was collected at the Biomedical Research Core Facilities Microscopy core, University of Michigan. This work was supported by NIH grant R01 AI071727 (to A.O.).

## References

1. Jouvenet N, Neil SJ, Bess C, Johnson MC, Virgen CA, Simon SM, Bieniasz PD. 2006. Plasma membrane is the site of productive HIV-1 particle assembly. PLoS Biol 4:e435.

2. Finzi A, Orthwein A, Mercier J, Cohen EA. 2007. Productive human immunodeficiency virus type 1 assembly takes place at the plasma membrane. J Virol 81:7476–90.

3. Adamson CS, Freed EO. 2007. Human immunodeficiency virus type 1 assembly, release, and maturation. Adv Pharmacol 55:347–87.

4. Bieniasz PD. 2009. The cell biology of HIV-1 virion genesis. Cell Host Microbe 5:550–8.

5. Freed EO. 2015. HIV-1 assembly, release and maturation. Nat Rev Microbiol 13:484–96.

6. Swanstrom R, Wills J. 1997. Synthesis, Assembly, and Processing of Viral Proteins. Retroviruses:p 263–334.

7. Sundquist WI, Kräusslich HG. 2012. HIV-1 assembly, budding, and maturation. Cold Spring Harb Perspect Med 2:a006924.

8. Garnier L, Bowzard JB, Wills JW. 1998. Recent advances and remaining problems in HIV assembly. AIDS 12 Suppl A:S5–16.

9. Balasubramaniam M, Freed EO. 2011. New insights into HIV assembly and trafficking. Physiology (Bethesda) 26:236–51.

10. Göttlinger HG, Sodroski JG, Haseltine WA. 1989. Role of capsid precursor processing and myristoylation in morphogenesis and infectivity of human immunodeficiency virus type 1. Proc Natl Acad Sci U S A 86:5781–5.

11. Bryant M, Ratner L. 1990. Myristoylation-dependent replication and assembly of human immunodeficiency virus 1. Proc Natl Acad Sci U S A 87:523–7.

12. Zhou W, Parent LJ, Wills JW, Resh MD. 1994. Identification of a membrane-binding domain within the amino-terminal region of human immunodeficiency virus type 1 Gag protein which interacts with acidic phospholipids. J Virol 68:2556–69.

13. Yuan X, Yu X, Lee TH, Essex M. 1993. Mutations in the N-terminal region of human immunodeficiency virus type 1 matrix protein block intracellular transport of the Gag precursor. J Virol 67:6387–94.

14. Ono A, Orenstein JM, Freed EO. 2000. Role of the Gag matrix domain in targeting human immunodeficiency virus type 1 assembly. J Virol 74:2855–66.

15. Ono A, Ablan SD, Lockett SJ, Nagashima K, Freed EO. 2004. Phosphatidylinositol (4,5) bisphosphate regulates HIV-1 Gag targeting to the plasma membrane. Proc Natl Acad Sci U S A 101:14889–94.

16. Freed EO, Orenstein JM, Buckler-White AJ, Martin MA. 1994. Single amino acid changes in the human immunodeficiency virus type 1 matrix protein block virus particle production. J Virol 68:5311–20.

17. Ono A, Freed EO. 1999. Binding of human immunodeficiency virus type 1 Gag to membrane: role of the matrix amino terminus. J Virol 73:4136–44.

18. Paillart JC, Göttlinger HG. 1999. Opposing effects of human immunodeficiency virus type 1 matrix mutations support a myristyl switch model of gag membrane targeting. J Virol 73:2604–12.

19. Spearman P, Horton R, Ratner L, Kuli-Zade I. 1997. Membrane binding of human immunodeficiency virus type 1 matrix protein in vivo supports a conformational myristyl switch mechanism. J Virol 71:6582–92.

20. Saad JS, Miller J, Tai J, Kim A, Ghanam RH, Summers MF. 2006. Structural basis for targeting HIV-1 Gag proteins to the plasma membrane for virus assembly. Proc Natl Acad Sci U S A 103:11364–9.

21. Saad JS, Loeliger E, Luncsford P, Liriano M, Tai J, Kim A, Miller J, Joshi A, Freed EO, Summers MF. 2007. Point mutations in the HIV-1 matrix protein turn off the myristyl switch. J Mol Biol 366:574–85.

22. Tang C, Loeliger E, Luncsford P, Kinde I, Beckett D, Summers MF. 2004. Entropic switch regulates myristate exposure in the HIV-1 matrix protein. Proc Natl Acad Sci U S A 101:517–22.

23. Zhou W, Resh MD. 1996. Differential membrane binding of the human immunodeficiency virus type 1 matrix protein. J Virol 70:8540–8.

24. Hermida-Matsumoto L, Resh MD. 1999. Human immunodeficiency virus type 1 protease triggers a myristoyl switch that modulates membrane binding of Pr55(gag) and p17MA. J Virol 73:1902–8.

25. Hill CP, Worthylake D, Bancroft DP, Christensen AM, Sundquist WI. 1996. Crystal structures of the trimeric human immunodeficiency virus type 1 matrix protein: implications for membrane association and assembly. Proc Natl Acad Sci U S A 93:3099–104.

26. Chukkapalli V, Hogue IB, Boyko V, Hu WS, Ono A. 2008. Interaction between the human immunodeficiency virus type 1 Gag matrix domain and phosphatidylinositol-(4,5)-bisphosphate is essential for efficient gag membrane binding. J Virol 82:2405–17.

27. Chan R, Uchil PD, Jin J, Shui G, Ott DE, Mothes W, Wenk MR. 2008. Retroviruses human immunodeficiency virus and murine leukemia virus are enriched in phosphoinositides. J Virol 82:11228–38.

28. Inlora J, Chukkapalli V, Derse D, Ono A. 2011. Gag localization and virus-like particle release mediated by the matrix domain of human T-lymphotropic virus type 1 Gag are less dependent on phosphatidylinositol-(4,5)-bisphosphate than those mediated by the matrix domain of HIV-1 Gag. J Virol 85:3802–10.

29. Inlora J, Collins DR, Trubin ME, Chung JY, Ono A. 2014. Membrane binding and subcellular localization of retroviral Gag proteins are differentially regulated by MA interactions with phosphatidylinositol-(4,5)-bisphosphate and RNA. MBio 5:e02202.

30. Alfadhli A, Barklis RL, Barklis E. 2009. HIV-1 matrix organizes as a hexamer of trimers on membranes containing phosphatidylinositol-(4,5)-bisphosphate. Virology 387:466–72.

31. Anraku K, Fukuda R, Takamune N, Misumi S, Okamoto Y, Otsuka M, Fujita M. 2010. Highly sensitive analysis of the interaction between HIV-1 Gag and phosphoinositide derivatives based on surface plasmon resonance. Biochemistry 49:5109–16.

32. Chukkapalli V, Oh SJ, Ono A. 2010. Opposing mechanisms involving RNA and lipids regulate HIV-1 Gag membrane binding through the highly basic region of the matrix domain. Proc Natl Acad Sci U S A 107:1600–5.

33. Shkriabai N, Datta SA, Zhao Z, Hess S, Rein A, Kvaratskhelia M. 2006. Interactions of HIV-1 Gag with assembly cofactors. Biochemistry 45:4077–83.

34. Chukkapalli V, Ono A. 2011. Molecular determinants that regulate plasma membrane association of HIV-1 Gag. J Mol Biol 410:512–24.

35. Purohit P, Dupont S, Stevenson M, Green MR. 2001. Sequence-specific interaction between HIV-1 matrix protein and viral genomic RNA revealed by in vitro genetic selection. RNA 7:576–84.

36. Hearps AC, Wagstaff KM, Piller SC, Jans DA. 2008. The N-terminal basic domain of the HIV-1 matrix protein does not contain a conventional nuclear localization sequence but is required for DNA binding and protein self-association. Biochemistry 47:2199–210.

37. Burniston MT, Cimarelli A, Colgan J, Curtis SP, Luban J. 1999. Human immunodeficiency virus type 1 Gag polyprotein multimerization requires the nucleocapsid domain and RNA and is promoted by the capsid-dimer interface and the basic region of matrix protein. J Virol 73:8527–40.

38. Chang CY, Chang YF, Wang SM, Tseng YT, Huang KJ, Wang CT. 2008. HIV-1 matrix protein repositioning in nucleocapsid region fails to confer virus-like particle assembly. Virology 378:97–104.

39. Cimarelli A, Luban J. 1999. Translation elongation factor 1-alpha interacts specifically with the human immunodeficiency virus type 1 Gag polyprotein. J Virol 73:5388–401.

40. Ott DE, Coren LV, Gagliardi TD. 2005. Redundant roles for nucleocapsid and matrix RNA-binding sequences in human immunodeficiency virus type 1 assembly. J Virol 79:13839–47.

41. Webb JA, Jones CP, Parent LJ, Rouzina I, Musier-Forsyth K. 2013. Distinct binding interactions of HIV-1 Gag to Psi and non-Psi RNAs: implications for viral genomic RNA packaging. RNA 19:1078–88.

42. Ramalingam D, Duclair S, Datta SA, Ellington A, Rein A, Prasad VR. 2011. RNA aptamers directed to human immunodeficiency virus type 1 Gag polyprotein bind to the matrix and nucleocapsid domains and inhibit virus production. J Virol 85:305–14.

43. Lochrie MA, Waugh S, Pratt DG, Clever J, Parslow TG, Polisky B. 1997. In vitro selection of RNAs that bind to the human immunodeficiency virus type-1 gag polyprotein. Nucleic Acids Res 25:2902–10.

44. Alfadhli A, Still A, Barklis E. 2009. Analysis of human immunodeficiency virus type 1 matrix binding to membranes and nucleic acids. J Virol 83:12196–203.

45. Alfadhli A, McNett H, Tsagli S, Bächinger HP, Peyton DH, Barklis E. 2011. HIV-1 matrix protein binding to RNA. J Mol Biol 410:653–66.

46. Chukkapalli V, Inlora J, Todd GC, Ono A. 2013. Evidence in support of RNA-mediated inhibition of phosphatidylserine-dependent HIV-1 Gag membrane binding in cells. J Virol 87:7155–9.

47. Kutluay SB, Zang T, Blanco-Melo D, Powell C, Jannain D, Errando M, Bieniasz PD. 2014. Global changes in the RNA binding specificity of HIV-1 gag regulate virion genesis. Cell 159:1096–1109.

48. Dick RA, Kamynina E, Vogt VM. 2013. Effect of multimerization on membrane association of Rous sarcoma virus and HIV-1 matrix domain proteins. J Virol 87:13598–608.

49. Mercredi PY, Bucca N, Loeliger B, Gaines CR, Mehta M, Bhargava P, Tedbury PR, Charlier L, Floquet N, Muriaux D, Favard C, Sanders CR, Freed EO, Marchant J, Summers MF. 2016. Structural and Molecular Determinants of Membrane Binding by the HIV-1 Matrix Protein. J Mol Biol 428:1637–55.

50. Dick RA, Vogt VM. 2014. Membrane interaction of retroviral Gag proteins. Front Microbiol 5:187.

51. Leventis PA, Grinstein S. 2010. The distribution and function of phosphatidylserine in cellular membranes. Annu Rev Biophys 39:407–27.

52. Llewellyn GN, Grover JR, Olety B, Ono A. 2013. HIV-1 Gag associates with specific uropod-directed microdomains in a manner dependent on its MA highly basic region. J Virol 87:6441–54.

53. Hafner M, Landthaler M, Burger L, Khorshid M, Hausser J, Berninger P, Rothballer A, Ascano M, Jungkamp AC, Munschauer M, Ulrich A, Wardle GS, Dewell S, Zavolan M, Tuschl T. 2010. PAR-CliP--a method to identify transcriptome-wide the binding sites of RNA binding proteins. J Vis Exp.

54. Yeung T, Gilbert GE, Shi J, Silvius J, Kapus A, Grinstein S. 2008. Membrane phosphatidylserine regulates surface charge and protein localization. Science 319:210–3.

55. Klein KC, Reed JC, Tanaka M, Nguyen VT, Giri S, Lingappa JR. 2011. HIV Gag-leucine zipper chimeras form ABCE1-containing intermediates and RNase-resistant immature capsids similar to those formed by wild-type HIV-1 Gag. J Virol 85:7419–35.

56. Li H, Dou J, Ding L, Spearman P. 2007. Myristoylation is required for human immunodeficiency virus type 1 Gag-Gag multimerization in mammalian cells. J Virol 81:12899–910.

57. Accola MA, Strack B, Göttlinger HG. 2000. Efficient particle production by minimal Gag constructs which retain the carboxy-terminal domain of human immunodeficiency virus type 1 capsid-p2 and a late assembly domain. J Virol 74:5395–402.

58. Crist RM, Datta SA, Stephen AG, Soheilian F, Mirro J, Fisher RJ, Nagashima K, Rein A. 2009. Assembly properties of human immunodeficiency virus type 1 Gag-leucine zipper chimeras: implications for retrovirus assembly. J Virol 83:2216–25.

59. Zhang Y, Qian H, Love Z, Barklis E. 1998. Analysis of the assembly function of the human immunodeficiency virus type 1 gag protein nucleocapsid domain. J Virol 72:1782–9.

60. Stauffer S, Rahman SA, de Marco A, Carlson LA, Glass B, Oberwinkler H, Herold N, Briggs JAG, Müller B, Grünewald K, Kräusslich HG. 2014. The Nucleocapsid Domain of Gag Is Dispensable for Actin Incorporation into HIV-1 and for Association of Viral Budding Sites with Cortical F-Actin, p 7893–903, J Virol, vol 88.

61. HIV Sequence Compendium 2018 Foley B LT, Apetrei C, Hahn B, Mizrachi I, Mullins J, Rambaut A, Wolinsky S, and Korber B, Eds. Published by Theoretical Biology and Biophysics Group, Los Alamos National Laboratory, NM, LA-UR 18-25673. HIV Sequence Compendium 2018.

62. Calnan BJ, Tidor B, Biancalana S, Hudson D, Frankel AD. 1991. Arginine-mediated RNA recognition: the arginine fork. Science 252:1167–71.

63. Li L, Vorobyov I, Allen TW. 2013. The different interactions of lysine and arginine side chains with lipid membranes. J Phys Chem B 117:11906–20.

64. Sokalingam S, Raghunathan G, Soundrarajan N, Lee SG. 2012. A study on the effect of surface lysine to arginine mutagenesis on protein stability and structure using green fluorescent protein. PLoS One 7:e40410.

65. Wu Z, Cui Q, Yethiraj A. 2013. Why do arginine and lysine organize lipids differently? Insights from coarse-grained and atomistic simulations. J Phys Chem B 117:12145–56.

66. Gaines CR, Tkacik E, Rivera-Oven A, Somani P, Achimovich A, Alabi T, Zhu A, Getachew N, Yang AL, McDonough M, Hawkins T, Spadaro Z, Summers MF. 2018. HIV-1 Matrix Protein Interactions with tRNA: Implications for Membrane Targeting. J Mol Biol 430:2113–2127.

67. Watanabe SM, Medina GN, Eastep GN, Ghanam RH, Vlach J, Saad JS, Carter CA. 2018. The matrix domain of the Gag protein from avian sarcoma virus contains a PI(4,5)P. J Biol Chem 293:18841–18853.

68. Eichorst JP, Chen Y, Mueller JD, Mansky LM. 2018. Distinct Pathway of Human T-Cell Leukemia Virus Type 1 Gag Punctum Biogenesis Provides New Insights into Enveloped Virus Assembly. MBio 9.

69. Ivanchenko S, Godinez WJ, Lampe M, Kräusslich HG, Eils R, Rohr K, Bräuchle C, Müller B, Lamb DC. 2009. Dynamics of HIV-1 Assembly and Release, PLoS Pathog, vol 5.

70. McMahon HT, Boucrot E. 2015. Membrane curvature at a glance. J Cell Sci 128:1065–70.

71. Dick RA, Goh SL, Feigenson GW, Vogt VM. 2012. HIV-1 Gag protein can sense the cholesterol and acyl chain environment in model membranes. Proc Natl Acad Sci U S A 109:18761–6.

72. Olety B, Veatch SL, Ono A. 2015. Phosphatidylinositol-(4,5)-Bisphosphate Acyl Chains Differentiate Membrane Binding of HIV-1 Gag from That of the Phospholipase Cδ1 Pleckstrin Homology Domain. J Virol 89:7861–73.

73. Mücksch F, Laketa V, Müller B, Schultz C, Kräusslich HG. 2017. Synchronized HIV assembly by tunable PIP. Elife 6.

74. Hogue IB, Grover JR, Soheilian F, Nagashima K, Ono A. 2011. Gag induces the coalescence of clustered lipid rafts and tetraspanin-enriched microdomains at HIV-1 assembly sites on the plasma membrane. J Virol 85:9749–66.

75. Ono A, Freed EO. 2001. Plasma membrane rafts play a critical role in HIV-1 assembly and release. Proc Natl Acad Sci U S A 98:13925–30.

76. Hogue IB, Hoppe A, Ono A. 2009. Quantitative fluorescence resonance energy transfer microscopy analysis of the human immunodeficiency virus type 1 Gag-Gag interaction: relative contributions of the CA and NC domains and membrane binding. J Virol 83:7322–36.

77. Kisseleva MV, Wilson MP, Majerus PW. 2000. The isolation and characterization of a cDNA encoding phospholipid-specific inositol polyphosphate 5-phosphatase. J Biol Chem 275:20110–6.

78. Nguyen KL, llano M, Akari H, Miyagi E, Poeschla EM, Strebel K, Bour S. 2004. Codon optimization of the HIV-1 vpu and vif genes stabilizes their mRNA and allows for highly efficient Rev-independent expression. Virology 319:163–75.

